# CTDPathSim: Cell line-tumor deconvoluted pathway-based similarity in the context of precision medicine in cancer ^*^

**DOI:** 10.1101/2020.06.13.149666

**Authors:** Banabithi Bose, Serdar Bozdag

**Affiliations:** Department of Mathematical and Statistical Sciences, Marquette University, Milwaukee Wisconsin USA; Department of Computer Science Marquette University Milwaukee Wisconsin USA

**Author notes:** Supplementary material of referred in this manuscript can be accessed at the pre-print version on bioRxiv and on Github page at https://github.com/bozdaglab/CTDPathSim under Creative Commons Attribution-NonCommercial 4.0 International Public License. Permission to make digital or hard copies of part or all of this work for personal or classroom use is granted without fee provided that copies are not made or distributed for profit or commercial advantage and that copies bear this notice and the full citation on the first page. Copyrights for third-party components of this work must be honored. For all other uses, contact the owner/author(s).

**Keywords:** Cell lines, Cancer Drugs, DNA Methylation, Gene Expression, Deconvolution, DE genes, REACTOME, TCGA, CCLE

## Abstract

In cancer research and drug development, human tumor-derived cell lines are used as popular model for cancer patients to evaluate the biological functions of genes, drug efficacy, side-effects, and drug metabolism. Using these cell lines, the functional relationship between genes and drug response and prediction of drug response based on genomic and chemical features have been studied. Knowing the drug response on the real patients, however, is a more important and challenging task. To tackle this challenge, some studies integrate data from primary tumors and cancer cell lines to find associations between cell lines and tumors. These studies, however, do not integrate multi-omics datasets to their full extent. Also, several studies rely on a genome-wide correlation-based approach between cell lines and bulk tumor samples without considering the heterogeneous cell population in bulk tumors. To address these gaps, we developed a computational pipeline, CTDPathSim, a pathway activity-based approach to compute similarity between primary tumor samples and cell lines at genetic, genomic, and epigenetic levels integrating multi-omics datasets. We utilized a deconvolution method to get cell type-specific DNA methylation and gene expression profiles and computed deconvoluted methylation and expression profiles of tumor samples. We assessed CTDPathSim by applying on breast and ovarian cancer data in The Cancer Genome Atlas (TCGA) and cancer cell lines data in the Cancer Cell Line Encyclopedia (CCLE) databases. Our results showed that highly similar sample-cell line pairs have similar drug response compared to lowly similar pairs in several FDA-approved cancer drugs, such as Paclitaxel, Vinorelbine and Mitomycin-c. CTDPathSim outperformed state-of-the-art methods in recapitulating the known drug responses between samples and cell lines. Also, CTDPathSim selected higher number of significant cell lines belonging to the same cancer types than other methods. Furthermore, our aligned cell lines to samples were found to be clinical biomarkers for patients’ survival whereas unaligned cell lines were not. Our method could guide the selection of appropriate cell lines to be more intently serve as proxy of patient tumors and could direct the pre-clinical translation of drug testing into clinical platform towards the personalized therapies. Furthermore, this study could guide the new uses for old drugs and benefits the development of new drugs in cancer treatments.

**CCS CONCEPTS:**

- Computational biology
- Genomics
- Systems biology
- Bioinformatics
- Genetics

**ACM Reference format:** Banabithi Bose, Serdar Bozdag. 2020. CTDPathSim: Cell line-tumor deconvoluted pathway-based similarity in the context of precision medicine in cancer.

## 1. Introduction

Due to genetic, molecular, and environmental factors affecting the cancer biology, each cancer patient is unique. Studies shows that tumor advancement, local growth, distant metastasis and genomic and epigenomic characteristics used to differ hugely within individual tumors *i.e.* in individual patients [1], [2], [8]. Hence, people that share the similar histopathological kind of tumor and tumor stage, and furthermore get a similar drug treatment, could have tumors with completely unique characteristics, behaviors, and drug response patterns. Hence, traditional therapy “one drug fits for all” should be changed as “patient-specific drugs” *i.e.* “personalized drugs”.

With the advancement of high throughput technologies, many repositories of the genome, proteome, transcriptome, and epigenome datasets, called multiple “omes” or multi-omics are being accessible to the researchers around the world. Integration of these different multi-omics layers such as gene expression, DNA methylation, mutation, DNA copy number aberration and publicly available clinical information on large cohorts of cancer patients, opens up an opportunity to leverage our understanding of personalized behavior of different tumor patients and paves the path towards the personalized drug treatment. But using patient’s molecular data in response prediction is limited [3].

Currently, in pre-clinical trials of pharmacogenomics studies, cell lines i.e. human tumor-derived cells, are utilized to understand the response of cancer drugs [4]. Several studies demonstrated that cell lines reflect numerous parts of the multi-omics signatures found in essential tumors and proposed that the same can be utilized as an intermediary for portraying the reaction to drug treatments [5], [6]. Several studies integrated the cell lines genomic feature with the chemical features of drugs to predict the cell line’s drug responses *in silico* [7]-[9]. However, *in vitro* culture of cell lines in general cause more genomic modifications in cell lines than primary tumors and likely give rise to systematic differences in the cancer patients’ and cells’ gene expression profiles [10], [11]. Hence, despite the fact that these cell line model CTDPathSim: Cell line-tumor deconvoluted pathway-based similarity in the context of precision medicine in cancer frameworks are exceptionally valuable for initial drug screening, these cell line-derived drug responses are far from actual patient’s drug responses.

Bridging the gap between cancer cell lines and primary tumors is an important and challenging task in developing personalized drug treatment for individual patients. Several studies integrated multi-omics datasets from primary tumors and cancer cell lines to find associations between cell lines and tumors. Sandberg *et al.* [12] devised a similarity measure called tissue similarity index (TSI), based on singular value decomposition (SVD) method using gene expression measurement of primary tumors and cell lines. TSI measures the distance between the singular-value decomposed gene expression pattern of a cell line and the average expression pattern of a patient cohort representing a particular tumor type. Dancik *et al.* [13] introduced a novel alignment method named Spearman’s rank correlation classification method (SRCCM) that measures similarity between cancer cell lines and tumor samples based on gene expression profiles to find cancer cell lines as biomarkers to patient survival. Liu *et al.* [14] used a transcriptome correlation analysis (TC analysis) to correlate cell lines and samples and found a substantial genomic differences between breast cancer cell lines and metastatic breast cancer samples. They also identified cell lines that more closely resemble the different subtypes of metastatic breast cancer patients. In a recent study, Warren *et al.* [15] developed an unsupervised alignment method called Celligner that mapped gene expression of around 12,000 tumors to gene expression of around 1,000 cell lines. This study found alignment of the majority of cell lines with tumor samples of the same cancer type while revealed systematic differences in others.

All these studies relied on a genome-wide correlation-based approach between cell lines and bulk tumor tissues derived from patients’ tumor. Bulk tissue derived from a patient’s tumor consists of heterogeneous cell population [16]-[18], hence it is important to consider the effect of these multiple cell types while establishing relationship between a tumor sample and a cell line. There exist computational deconvolution methods, such as CIBERSORT [19] and TIMER [20] that deconvolve the cell type proportions using cell type-specific gene expression references. Also, there are deconvolution methods, such as MethylCIBERSORT [21], MeDeCom [22] and EDec [23], that deconvolute the cell type proportions from DNA methylation data of the bulk tumor tissue.

In the present study, we developed a computational pipeline, CTDPathSim, a pathway activity-based approach to compute similarity between tumor samples and cell lines at genetic, genomic, and epigenetic levels integrating multi-omics datasets, addressing bulk tumor heterogeneity for different cell type populations.

Several studies show the association of DNA methylation in blood with the immune response and inflammation in cancer[24]-[29]. Also, studies suggest that blood-based methylation profile can be able to capture the distribution of different cell types, especially for immune cell type profiles [30], [31]. In cancer, roles of immune cells are important and therapies targeting these immune cells are common [32]-[35]. Following these studies, we prepared a reference methylation profiles using different immune cell types from peripheral blood samples. We computed sample-specific deconvoluted expression and methylation profiles utilizing a deconvolution-based algorithm with our reference methylation profiles. We computed pathway-specific differentially expressed (DE) genes and differentially methylated (DM) genes and computed a similarity score using Spearman rank correlation between each sample-cell line pair using these genes. We assessed CTDPathSim using breast and ovarian cancer data from TCGA database and cancer cell lines from CCLE. CTDPathSim outperformed TSI, SRCCM, TC analysis and Celligner in predicting similarity scores that recapitulate the known drug responses between TCGA patients and CCLE cell lines. Also, CTDPathSim selected higher number of significant cell lines belonging to the same cancer types than other methods. Our aligned cell lines to samples were found to be clinical biomarkers for patients’ survival whereas unaligned cell lines were not. Furthermore, our high score sample-cell line pairs had higher number of significant overlapping biological pathways than low score sample-cell line pairs.

## 2. Materials and methods

### 2.1. CTDPathSim

We developed CTDPathSim, a computational pipeline to compute similarity scores between patient samples and cell lines using a pathway activity-based approach. CTDPathSim integrates sample-specific expression and methylation profiles utilizing a deconvolution method. CTDPathSim has four main computational steps (Fig. 1). In order to capture the accurate methylation signal, in the first step, we employed a deconvolution-based algorithm and inferred sample’s deconvoluted methylation profile with proportions of different cell types in a sample’s bulk tumor tissue. In the second step, using the estimated proportions of different cell types in the previous step, we used bulk tumor gene expression data to infer sample-specific deconvoluted expression profile. In the third step, we computed sample-specific and cell line-specific DE genes and DM genes among the same cancer type cohort. Then using sample-specific DE genes, we computed enriched biological pathways for each patient. Similarly, using cell line-specific DE genes, we computed cell line-specific enriched biological pathways. In the last step, we computed a union of pathways between each sample-cell line pair and computed a union of DE genes for each pair. We used DE genes that were enriched in union of pathways of each pair to compute the Spearman rank correlation between each sample-cell line pair using sample’s deconvoluted expression profile and cell line’s expression profile. Similarly, we found DM genes that were enriched in union of pathways of each pair. We computed the Spearman rank correlation using common DM genes in each sample-cell line pair using sample’s deconvoluted methylation profile and cell line’s methylation profile. The entire pipeline of CTDPathSim is illustrated in Fig. 1.

**Figure 1:**
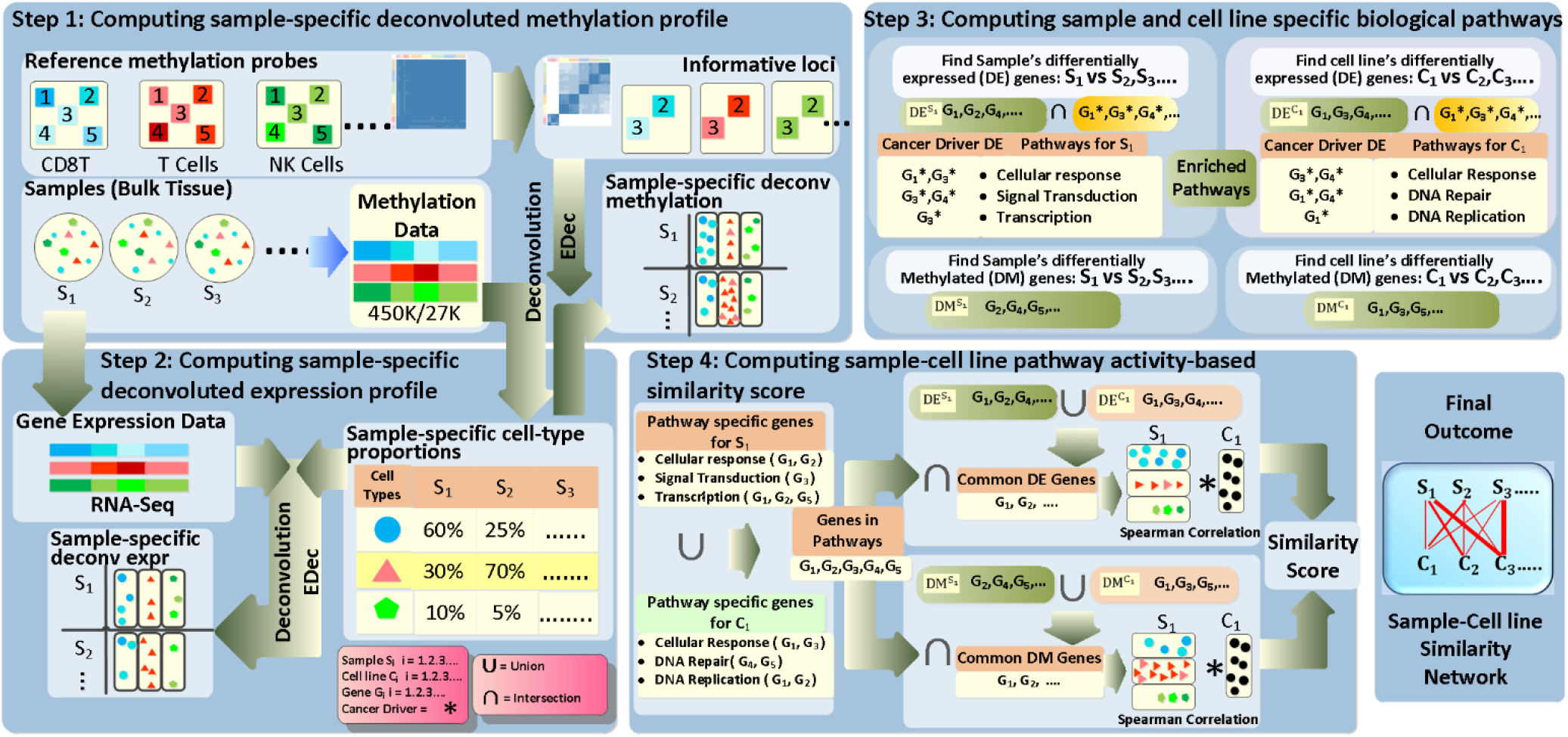
Flowchart of the CTDPathSim pipeline.

#### 2.1.1 Computing sample-specific deconvoluted methylation profile

In order to capture the accurate methylation signal, in the first step, we employed a deconvolution-based algorithm, EDec [23], based on a heuristic for constrained matrix factorization and inferred sample’s deconvoluted methylation profile with proportions of different cell types in sample’s bulk tumor tissue.

To run EDec, we prepared a reference methylation profile of different cell types from peripheral blood and used that to deconvolute methylation signal of tumor samples. Our reference file contained eight different cell types, such as, B cell, natural killer (NK), CD4T, CD8T, monocytes, neutrophils, adipocytes, cortical neurons and vascular endothelial with replicates. Using EDec, we identified a set of important loci in the references that are likely to exhibit variation in methylation levels across different constituent cell types of a given tumor sample in a feature selection manner. EDec identified constituent cell types by comparing their deconvoluted molecular profiles to the reference profiles. Using, the inferred proportion and probe centric methylation profile of constituent cell types, we computed the sample-specific methylation profiles for each tumor sample. We ran this step for 450K methylation probes and 27K methylation probes separately. At the end of this step, gene-specific beta values were calculated separately for both platforms. For the 450K platform, average beta value for promoter-specific probes were considered due to their role in transcriptional silencing [36]. Given lower coverage in the 27K platform, we utilized all the probes. In this case, we set the DNA methylation of a gene as the average of beta values of all its probes.

#### 2.1.2 Computing sample-specific deconvoluted expression profile

We computed corrected expression profile for each sample using deconvolution. Firstly, we computed cell typespecific expression profiles for our reference immune cells. For this purpose, we used computed cell-type proportions for each sample in previous step and gene expression data of bulk tumor samples. We applied EDec stage 2 to compute cell-type specific expression profiles. Then, using cell type-specific proportions and cell type-specific expression, we inferred the sample-specific deconvoluted expression profile of each tumor sample.

#### 2.1.3 Computing sample and cell line-specific biological pathways

In this step, we computed enriched biological pathways listed in REACTOME [37] database for each sample and cell line. To find these pathways, we computed sample-specific DE genes and sample-specific DM genes among the same cancer type cohort. We considered gene expression data and gene centric methylation data with beta values [38] with bulk tumor data for differential analysis. To compute sample-specific DE genes, we computed the median expression of a gene across all samples and computed fold change of that gene w.r.t the computed median expression. A gene was considered up if the fold change ≥ 4 and down if the fold change ≤ 0.25. To compute sample-specific DM genes, we computed M-values [39] from gene centric beta values and computed the median of gene centric M-values across all sample for each gene. We considered a gene hypermethylated if the M-value fold change ≥ 4 and hypomethylated if fold change ≤ 0.25. Likewise, using the same thresholds, we computed cell line-specific DE genes and DM genes in the same cancer type.

To identify enriched pathways for each sample, we used only the frequently mutated DE genes of that sample. We obtained the frequently mutated cancer driver genes from the COSMIC [40] database. Using these cancer driver DE genes, we identified enriched biological pathways listed in REACTOME via a pathway enrichment tool *Pathfinder* [41]. Similarly, we identified cancer driver DE genes for each cell line and computed cell line-specific enriched REACTOME based pathways. We considered the pathways that were significantly enriched with FDR corrected hypergeometric p-values < 0.05.

#### 2.1.4 Computing sample-cell line pathway activity-based similarity score

We computed Spearman rank correlation between each sample-cell line pair using expression and methylation profiles. We gathered all the genes that were present in the union of enriched REACTOME pathways of a sample-cell line pair. Then, using a union of sample and cell line’s DE genes, we found DE genes that were present in the gene list of the union of enriched pathways. We used these pathway-based DE genes to compute the Spearman rank correlation between each sample-cell line pair using sample’s deconvoluted expression profile and cell line’s expression profile. Similarly, using a union of sample and cell line’s DM genes, we found DM genes that were present in the gene list of the union of enriched REACTOME pathways. We used sample’s deconvoluted methylation profile and cell line’s methylation profile to compute Spearman rank correlation using pathway-based DM genes. We scaled expression-based and methylation-based correlations into 0 to 1 and then combined these two correlation scores to get a unified similarity score for each sample-cell line pair.

### 2.2. Running state-of-the-art methods

We compared CTDPathSim with four state-of-the-art methods, namely, TSI [18], SRCCM [19], TC analysis [20] and Celligner [21] by running them on BRCA and OV datasets from TCGA and cancer cell line datasets from CCLE.

#### 2.2.1 Running TSI method

For TSI method, we performed z-normalization of RNA-Seq data for each gene of samples and cell lines. We then applied SVD [18] on normalized gene expression matrix of samples. Then each sample was projected into SVD space by measuring its correlation to the Eigenarrays (16 for BRCA; 20 for OV, Supplemental Fig. 1) with the largest singular values that explain the most variance of the data. Similarly, we projected each cell line into SVD space. Then, we computed a Pearson correlation score between each patient and cell line as a similarity score, namely, TSI score. We applied z-normalization on TSI scores computed between 1,102 BRCA samples and 1,019 cell lines and between 374 OV samples and 1,019 cell lines.

#### 2.2.2 Running SRCCM

For SRCCM method, we used 43 breast cancer driver genes [40] that are considered to be related with different histologic grades among breast cancer patients [42] and used Spearman rank correlations as similarity scores, namely, SRCCM scores. We applied z-normalization on TSI score computed between 1,102 TCGA samples with 1,019 CCLE cell lines and between 374 OV samples with 1,019 cell lines.

#### 2.2.3 Running TC analysis

For TC analysis, we used 30,681 common genes between CCLE and TCGA RNA-Seq gene expression data with log2 transformation. Then, we rank-transformed gene RPKM values for each CCLE cell line and ranked all the genes according to their expression variation across all CCLE cell lines. Following the SRCCM method, the 1000 most variable genes were kept as “marker genes”. We used these “marker genes” to compute Spearman rank correlation, namely, TC scores, between each cell line and sample pair. We computed TC scores between 1,102 TCGA samples with 1,019 CCLE cell lines and between 374 OV samples with 1,019 cell lines.

#### 2.2.4 Running Celligner

For Celligner, we downloaded RNA-Seq data in TPM (transcript per million reads) of 12,236 TCGA samples and 1,249 CCLE cell lines as log2(TPM + 1) transformation from Cancer Dependency Map [https://depmap.org] data portal spanning over 37 different cancer types. We used cell line and TCGA gene expression data without ‘non-coding RNA’ and ‘pseudogene’ as input in Celligner pipeline. Celligner first identified and removed the expression signatures with excess ‘intra-cluster’ variance in the sample compared to cell line data. Then, using differentially expressed genes between clusters in the data it ran contrastive Principal Component Analysis (cPCA) followed by a batch correction method, MNN (mutual nearest neighbors) that aligned similar sample-cell line pairs to produce a corrected gene expression data. We further filtered this gene expression matrix with 1,088 BRCA and 372 OV samples from TCGA and 1,015 CCLE cell lines. Then, we computed 1,000 most variable genes in cell lines as “marker genes”. We used these “marker genes” with Celligner’s gene expression data to correlate each sample and cell line using a Spearman rank correlation, namely, Celligner score. We applied z-normalization on Celligner scores computed with BRCA and OV cohorts.

### 2.3. Datasets

#### 2.3.1 Data preprocessing for running CTDPathSim

We tested CTDPathSim on breast cancer (BRCA) and ovarian cancer (OV) samples from TCGA and cancer cell lines from CCLE database. Using *TCGAbiolinks* [43], we downloaded RNA sequencing (RNA-Seq) data in HTSeq-FPKM format. We downloaded DNA methylation data from Infinium HumanMethylation27 Bead-Chip (27K) and Infinium HumanMethylation450 Bead-Chip (450K) platforms. We downloaded CCLE_RNAseq_gene_rpkm _10180929.gct.gz file for gene expression and CCLE_RRBS_ cgi_CpG_clusters_20181119.txt.gz file for DNA methylation data of cell lines from CCLE database. Using R package *BioMethyl* [44], we removed CpG sites that have missing values in more than half of the samples and imputed the rest of missing values by integrating the *Enmix* [45] R package, with default parameters. From COSMIC (Catalogue Of Somatic Mutation In Cancer) [40] cancer gene census project, we downloaded a list of ~700 cancer driver genes that are found to be frequently mutated in different cancer types.

To prepare the reference methylation profiles for the deconvolution step, we processed raw methylation probes provided in *FlowSorted.Blood.450k* [46] R package that consists methylation probes from peripheral blood samples with five different cell types, such as, B cell, natural killer (NK), CD4T, CD8T, monocytes, generated from adult men with replicates. We also processed raw methylation data available at NCBI Gene Expression Omnibus (GEO) database repository with the dataset identifier GSE122126 for three more cell types such as, adipocytes(450K), cortical neurons and vascular endothelial cells with replicates. We processed these raw methylation files using R package *minfi* [47] to prepare our reference methylation probes of different cell types. Two different reference files for 450K probes and 27K probes with eight different cell types were prepared.

#### 2.3.2 Data preprocessing for evaluating CTDPathSim’s results

To assess the concordance of drug response between sample-cell line pairs and evaluating the prognostic relevance of cell lines as clinical biomarkers for patients’ survival, we considered four different survival endpoints, namely, Overall Survival (OS), Progression Free Survival (PFI), Disease Free Survival (DSS) and DFI (Disease Free Interval). In OS, patients who were dead from any cause considered as dead, otherwise censored. In PFI, patients having new tumor event whether it was a progression of disease, local recurrence, distant metastasis, new primary tumor event, or died with the cancer without new tumor event, including cases with a new tumor event whose type is N/A were considered as dead and all other patients were censored. DFI was similar to PFI with the inclusion of censored patients with new primary tumor in other organ; patients who were dead with tumor without new tumor event and patients with stage IV were excluded. In DSS, disease-specific survival time in days, last contact days or death days, whichever was larger was used to identify dead vs censored patients [48].

To compare the drug responses between sample-cell line pairs, we compiled our ground truth drug response data for patients from TCGA database using OS and PFI as clinical end points. We also considered the time intervals in which drugs were applied. We considered survival status 0 (i.e. censored), representative of sensitive drug response whereas survival status 1 (i.e. dead), representative of resistant drug response as we mapped patients’ survival status to drug response using the following criteria:

- *Drug response = OS status; if PFI status = 0 and PFI.time = OS.time*
- *Drug response = PFI status; if PFI = 1 and PFI.time = OS.time*
- *Drug response = PFI status; if drug application end days ≤ PFI days and PFI = 1 with PFI. time < OS. time*
- *Drug response = OS status; if drug application end days ≤ PFI days and PFI = 0 with PFI.time < OS.time*
- *Drug response = OS status; if drug application starts days ≥ PFI days*

For cell lines’ drug response data, we considered IC50 (the concentration of a drug that is required for 50% inhibition *in vitro*) values as drug response quantification. Furthermore, ln (IC50) < −2.0 was used to define positive (i.e., sensitive) anticancer drug response and other ln (IC50) values were considered as negative (i.e., resistant) drug response.

Since, common drugs between TCGA and CCLE were fewer, we used cancer cell lines drug response data with IC50 values from GDSC database. We found around 900 overlapping cell lines between CCLE and GDSC databases. We processed IC50 values of GDSC drug in similar way like we did for CCLE. We retrieved around 60 common drugs between TCGA patients and cell lines from CCLE and GDSC. We found 13 common drugs between 368 cell lines and 607 BRCA cohort and 14 common drugs between 368 cell lines and 307 OV patient cohort. A list of these drugs is provided in Supplemental Table 1.

Since the known drug screening data between TCGA samples and cell lines were limited, we included structurally similar drugs in our evaluation. We retrieved PaDEL descriptor [49] of drug’s chemical properties and 2D and 3D structural profiles of 378 drugs in GDSC using a line notation of simplified molecular-input line entry system (SMILES) [50] from PubChem [51] database. We computed structural similarities using Pearson correlation between different drugs from TCGA and GDSC.

### 2.4. Hypergeometric test for predicted similarity scores evaluating sample-cell lines’ known drug response

We evaluated CTDPathSim based on known drug responses between cancer patients and CCLE cell lines.

We computed the similarities scores such that we had discrete similarity thresholds, each with multiple sample-cell line pairs. For a specific drug, D, we computed the number of sample-cell line pairs for all similarity thresholds, namely, Set 1. Then, we computed the number of sample-cell line pairs for a specific similarity threshold, K, namely, Set 2. For drug, D, we considered the similarity thresholds that have significant overlap between Set 1 and Set 2 (hypergeometric p-value < 0.05) to evaluate drug response match and mismatch between different sample-cell line pairs.

## 3. Results

We developed a computational pipeline, CTDPathSim, a pathway activity-based approach to compute similarity between tumor samples and cell lines at genetic, genomic, and epigenetic levels.

We assessed CTDPathSim using breast invasive carcinoma (BRCA) and ovarian serous cystadenocarcinoma (OV) samples from TCGA with around 1,000 cell lines from CCLE spanning over 24 different types of cancer.

We downloaded DNA methylation data from Infinium HumanMethylation27 Bead-Chip (27K) and Infinium HumanMethylation450 Bead-Chip (450K) platforms. For BRCA we had 1,236 samples and for OV we had 623 samples with methylation profiles combined from both the platforms. We downloaded RNA-Seq data of 1,097 BRCA and 374 OV primary tumor samples. We retrieved 1,073 unique samples for BRCA and 352 samples for OV with both DNA methylation and gene expression data.

After running step 1 and step 2 (see Materials & methods *2.1.1* and *2.1.2*) of our pipeline, we had deconvoluted methylation and expression profiles of each BRCA sample based on three different immune cell types, namely cytotoxic T cell (CD4T), vascular endothelial cell and monocytes. In OV, deconvolution using 450K DNA methylation data resulted in three different cell types namely cortical neurons, vascular endothelial cell and adipocytes, whereas using 27K DNA methylation data resulted in monocytes in an addition to the other three cell types.

We downloaded expression data of 1,019 cell lines and methylation data of 831 cell lines from CCLE. There were 824 cell lines with both data types.

We identified enriched biological pathways from REACTOME database using a group of cancer driver (mutated) genes that were present in patient’s DE genes and cell line’s DE genes (see Materials & methods *2.1.3*). We provided the boxplots of the counts of DE genes, DM genes in Supplemental Fig. 2 and a list of REACTOME pathways for patients and cell lines in Supplemental Table 2.

In the final step of CTDPathSim, we computed similarity between BRCA/OV samples and 1,108 cell lines that had enriched biological pathways (see Materials & methods *2.1.4*). We excluded one cell line that had no enriched pathway from REACTOME.

In our final step, for BRCA, we computed Spearman correlation between methylation profiles of 1,073 samples and 824 cell lines, and between expression profiles of 1,073 samples and 1,018 cell lines. We scaled Spearman similarities to the range of 0 to 1 and computed an average similarity score taking a mean of expression-based correlation and methylation-based correlation for each cell line-sample pair. Since we did not have DNA methylation data for 194 cell lines, there were sample-cell line pairs that had no methylation-based scores but only expression-based scores. For these pairs, we considered only expression-based score in our final similarity score. We applied z-normalization on our final scores such that the mean score was 0 with standard deviation 1 (Fig. 2A). Similarly, for ovarian cancer, we computed final similarity scores between 352 samples and 1,018 cell lines (Fig. 2B).

**Figure 2:**
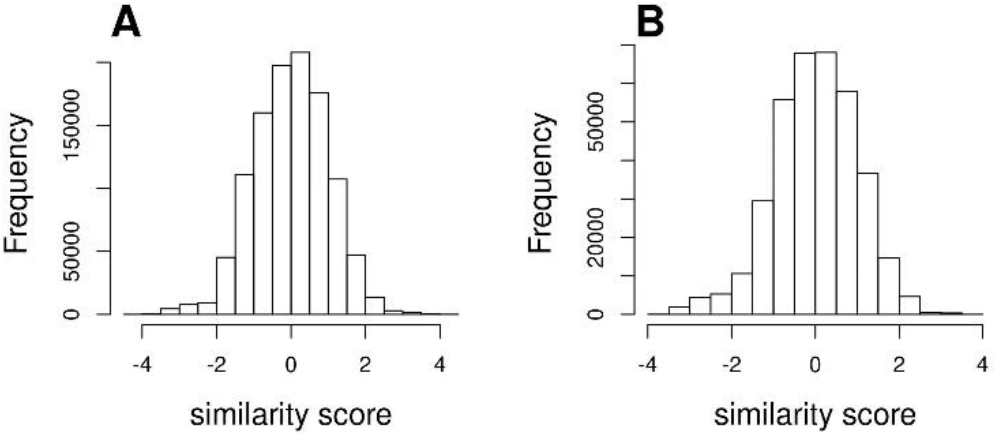
Histograms of sample-cell line pairwise similarity scores predicted by CTDPathSim. **A)** z-normalized similarity scores with 1,073 BRCA samples and 1,018 CCLE cell lines; **B)** z-normalized similarity scores with 352 OV samples and 1,018 CCLE cell lines.

## 4. Evaluation of CTDPathSim’s results

We performed several tests to assess the similarity scores of sample-cell line pairs computed by CTDPathSim. CTDPathSim outperformed the state-of-the-art methods by inferring high similarity score for sample-cell line pairs that belong to the same cancer type. CTDPathSim computed the similarity scores that were found to be significantly associated with the responses of known cancer drugs (see Materials & methods *2.3.2*) between CCLE cell lines and TCGA samples. We found that cell lines with high CTDPathSim scores i.e. aligned cell lines were the clinical biomarkers for patients’ survival whereas unaligned cell lines were not. Furthermore, CTDPathSim computed aligned sample-cell line pairs that had higher number of significant overlapping biological pathways compared to unaligned sample-cell line pairs. In the following sections we presented the details of the assessment of our tool, CTDPathSim

### 4.1. CTDPathSim outperformed four existing methods in selection of significant cell lines belonging to the same cancer type

We compared CTDPathSim with the four state-of-the-art methods (See Section 2.2), namely, TSI [18], SRCCM [19], TC analysis [20] and Celligner [21] with RNA-Seq data of BRCA and OV samples from TCGA and cancer cell lines from CCLE. For all the methods, we evaluated the enrichment of breast cancer cell lines in BRCA samples, and ovarian cancer cell lines in OV samples. We used a high similarity threshold (≥ 1.5) and a low similarity threshold (≤ −1.5) for the sample-cell line pairs for all the methods, except, TSI. For TSI, we used low similarity ≤ −1 to include sufficient number of sample-cell line pairs to run this analysis. In these thresholds, we computed highly and lowly similar cell lines for each sample and checked if the cell lines for each BRCA sample had highly overlapping breast cancer cell lines. Similarly, we checked if the cell lines for each OV sample had highly overlapping ovarian cancer cell lines. We found that, CTDPathSim was able to select breast cancer cell lines for BRCA samples (Fig. 3A1) and ovarian cancer cell lines for OV samples (Fig. 3B1) with significant hypergeometric p-values (< 0.05) in high similarity thresholds compared to low similarity thresholds. On the other hand, TSI method (Fig. 3A2, 3B2), TC analysis ((Fig. 3A3, 3B3) and Celligner (Fig. 3A4, 3B4), produced highly overlapping p-values between high and low similarity thresholds, whereas SRCCM performed little better than these three comparing methods for both BRCA (Fig. 3A5) and OV (Fig. 3B5).

**Figure 3:**
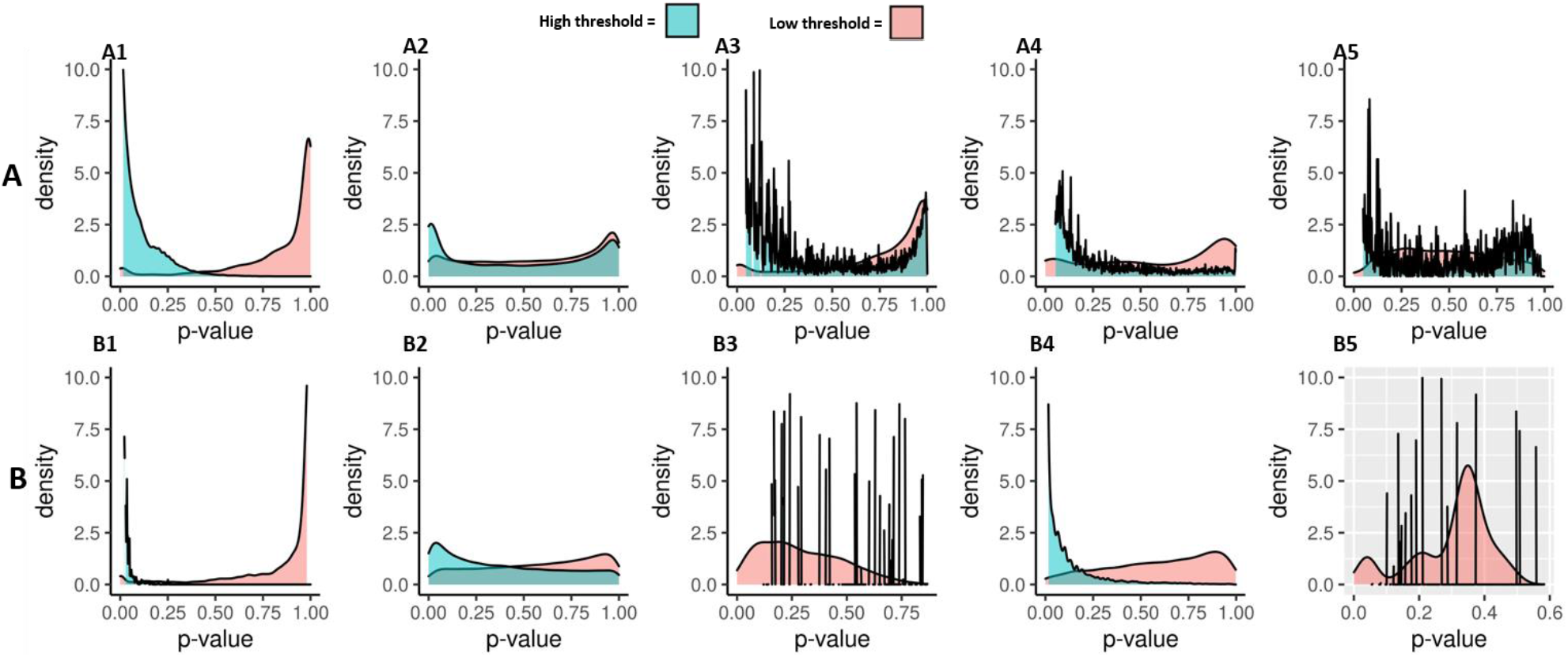
Density plots of hypergeometric p-values for lineage specific cancer cell lines. **A)** density plots of p-values for breast cancer cell lines’ enrichment in BRCA samples: A1) CTDPathSim; A2) TSI method; A3) TC analysis; A4) SRCCM; A5) Celligner; **B)** density plots of p-values for ovarian cancer cell lines’ enrichment in OV samples: B1) CTDPathSim; B2) TSI method; B3) TC analysis; B4) SRCCM; B5) Celligner. We used “high (≥ 1.5) and (low ≤-1.5) similarity thresholds for all methods (only in TSI, we used low similarity ≤ −1 to include sufficient number of sample-cell line pairs to run this analysis) in both cancer types.

### 4.2. CTDPathSim outperformed four existing methods for predicting similarity scores that recapitulate the known drug responses between tumor samples and cell lines

We compared the computed similarity scores of CTDPathSim with the scores computed by TSI, SRCCM, TC analysis and Celligner to check whether these scores were aligned with the known drug responses between TCGA BRCA and CCLE cell lines. We used z-normalized scores of all five methods and compared the known drug responses of 13 common drugs between 382 cell lines and 681 BRCA samples. Among these 13 drugs, we considered only those drugs that were filtered for class imbalance of matching and mismatch drug response information in our ground truth data (See Materials & methods *2.3.2*). For this purpose, we computed an imbalance ratio (i.e. size of minority class/size of majority class) for each drug with R package *imbalance* [52]. Since lower imbalance ratio shows more imbalance in matching and mismatching cases between sample-cell line pairs for drug response of a particular drug, we eliminated the drugs from this analysis that were below 0.2 imbalance ratio. Then, for each drug, we computed hypergeometric p-values for different similarity scores considering the number of sample-cell line pairs that had known drug response in our ground truth data (See Materials & methods 2.4). We considered only those scores that had significant p-value (< 0.05) as eligible scores for checking drug response. Furthermore, we considered the scores for which we have at least 10 percent difference between matching and mismatch drug response information to decide whether these scores had cumulative i.e. overall matching or mismatch drug response.

We found that, there were higher number of drug response mismatch between sample and cell lines in low CTDPathSim scores (Fig 4B1), whereas, there were higher number of matching incidents in higher similarities (Fig 4C1). Majority of the drugs compared with our predicted scores, showed trends of concordance in drug response with CTDPathSim scores i.e. in low similarities, matching drug response between sample-cell line pairs were lower and as our similarity score became higher the matching drug response percentage was improved with significant p-value (<0.05) (Fig 4B1, 4C1). On the other hand, all other four methods were not able to capture the trends of the known drug responses with their predicted scores (Fig 4B2-B5; Fig 4C2-C5). In Fig 4A, we showed the drug response concordance with sample-cell line similarity scores of Vinorelbine between BRCA samples and CCLE cell lines with computed similarity scores by five different methods. The matching percentage of the response of drug Vinorelbine increases as the similarity score increases only in CTDPathSim (Fig 4A1).

**Figure 4:**
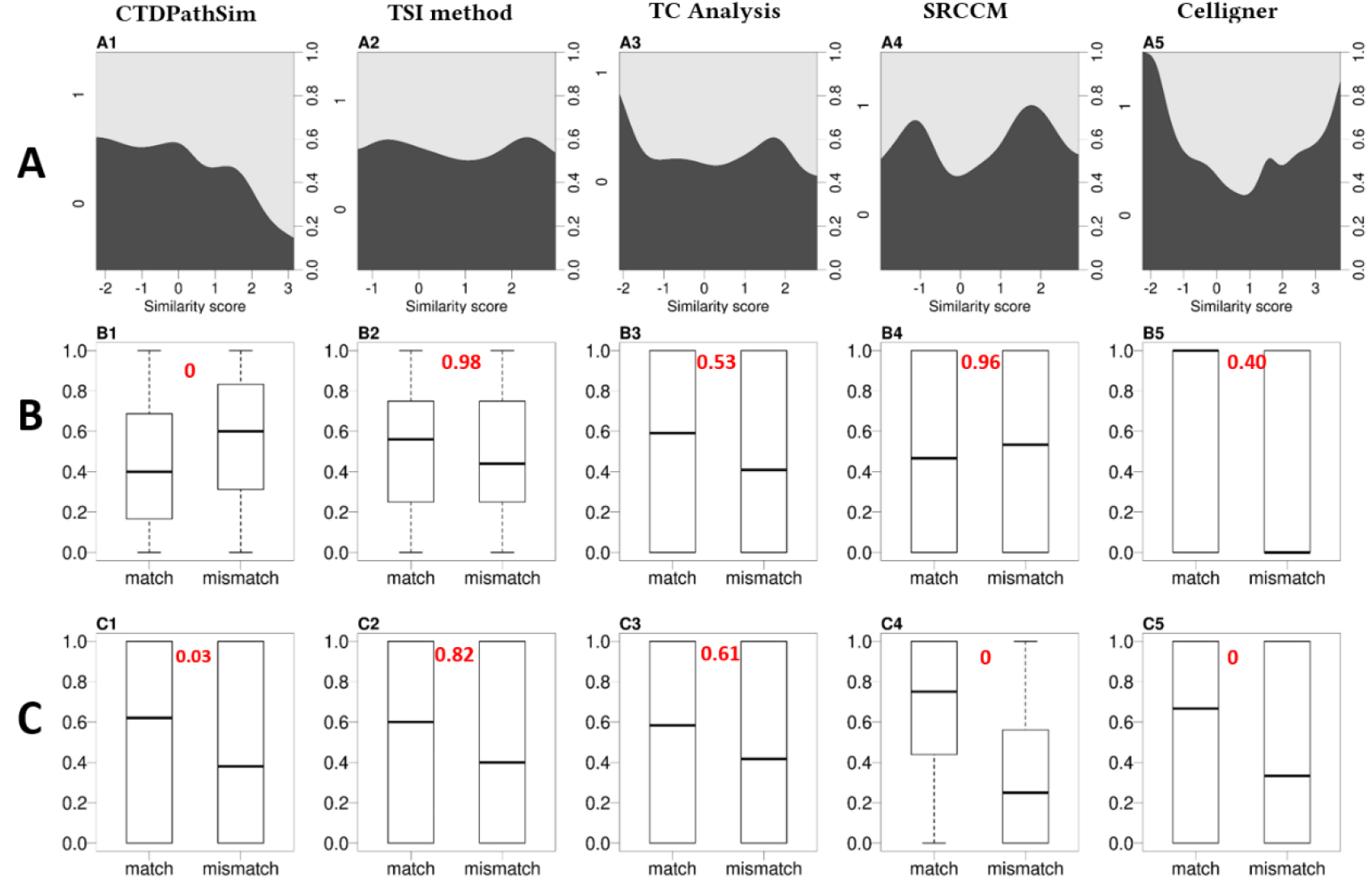
Drug response concordance with sample-cell line similarity scores. **A)** Conditional density plot of matching (1 ~ grey) and mismatch (0 ~ black) of Vinorelbine’s response between BRCA samples and CCLE cell lines with computed similarity scores by five different methods: A1) CTDPathSim, A2) TSI method, A3) TC analysis, A4) SRCCM, A5) Celligner. The matching percentage of response of drug vinorelbine increases as the similarity score increases in A1; whereas in A2-A5, this drug does not show concordance between response and similarity scores. **B)** Boxplots of matching and mismatch response percentage with common drugs between TCGA BRCA cohort and CCLE cell lines in a low similarity threshold (≤ −1.5) for all the methods except TSI threshold (≤ −1) (with a Wilcoxson p-value in each box in red: B1) CTDPathSim shows lower median value of matching drug response in low similarity threshold with significant p-value (<0.05), whereas all other four methods in B2-B5, there was no difference between medians of matching and mismatch responses (p-values > 0.05). **C)** Boxplots of matching and mismatch response percentage with common drugs between TCGA BRCA cohort and CCLE cell lines in a high similarity threshold (≥ 1.5)for the five methods with a Wilcoxson p-value in each box in red: C1) CTDPathSim shows higher median value of matching drug response in high similarity threshold with significant p-value (<0.05), whereas for TSI method and TC analysis in C2 and C3, there was no difference between medians of matching and mismatch responses ( p-values > 0.05); C4)SRCCM and C5)Celligner also show significant higher matching response in high similarity threshold.

We provided the conditional density plots for drug response vs similarity scores of all these drugs for different methods in Supplemental Fig. 3-8. Table 1 shows the total number of drugs that were used to test different methods after applying our filtering criteria with percentages of aligned drugs i.e. the drugs that had good concordance with similarity scores and unaligned drugs *i.e.* the drugs that had poor concordance with similarity scores. We also listed the percentage of the drugs, for which these different methods were not able to identify aligned or unaligned drug response trend based on conditional density plots, as undecided drugs.

**Table 1:** Number of drugs that were used by different methods with aligned, unaligned, and undecided drug response trends with similarity scores predicted by different methods for BRCA cohort and CCLE cell lines

| Method | No. of selected drugs | Aligned | Unaligned | Undecided |
| --- | --- | --- | --- | --- |
| CTDPathSim | 7 | 40% | 10% | 40% |
| SVD method | 6 | 30% | 50% | 20% |
| TC analysis | 5 | 40% | 40% | 20% |
| SRCCM | 7 | 30% | 30% | 40% |
| Celligner | 6 | 20% | 60% | 20% |

### 4.3. Aligned cell lines to samples were found to be clinical biomarkers for patients’ survival

We checked whether our aligned cell lines to BRCA samples could be considered as clinical biomarkers for patients. We considered sample-cell line pairs with z-normalized CTDPathSim scores with high similarity threshold (≥ 3) and gathered all the cell lines for each sample in this group. We took an average gene expression of these aligned cell lines as proxy of sample’s gene expression and ran a multivariate Cox regression [49] with proxy expression of each gene using other clinical variables of BRCA cohort, such as age, race, histology, HER2 status, estrogen status and progesterone status as independent variables. We considered four different survival end points, Overall Survival (OS), Progression Free Survival (PFI), Disease Free Survival (DSS) and DFI (Disease Free Interval) (see Materials & methods *2.3.2*). The genes with Cox regression p-value < 0.05 were considered as prognostic genes.

We computed Hazard ratios (HR) associated with these prognostic genes as proxy of samples’ gene and called these HR as cell-HR. For this same samples group, we ran the multivariate Cox regression with the actual gene expression of patient samples with the same clinical variables and four survival end points and computed HR associated with prognostic genes, namely, patient-HR. Then, we computed a Spearman rank correlation score between cell-HR and patient-HR using overlapping prognostic genes. We performed the similar Cox regression in sample-cell lines group with low similarity threshold (≤ −2) and computed Spearman correlation of cell-HR and patient-HR using overlapping prognostic genes in this group.

Table 2 shows that Spearman correlation scores computed for highly similar sample-cell lines group were higher in OS, PFI and DSS along with significant p values (< 0.05) in PFI and DSS, whereas, low similarity group had slightly higher score only with DFI, however the p-value was not significant (>0.05). Furthermore, the number of overlapping prognostic genes were higher in highly similar group than in lowly similar group.

**Table 2:** Correlation of hazard ratios between cell lines and samples using prognostic genes in high and low similarity thresholds predicted by CTDPathSim with BRCA cohort. Correlation with significant p-values are with asterisk (*) symbol.

| | Similarity $\geq 3$ | | | | Similarity $\leq -2$ | | | |
| --- | --- | --- | --- | --- | --- | --- | --- | --- |
| Survival end points | OS | PFI | DFI | DSS | OS | PFI | DFI | DSS |
| Prognostic genes with cell lines | 287 | 644 | 1772 | 961 | 2373 | 145 | 27 | 397 |
| Prognostic genes with patients | 2039 | 1893 | 1615 | 2159 | 2344 | 2110 | 1605 | 2217 |
| Overlapping Prognostic genes | 17 | 45 | 103 | 46 | 194 | 9 | 4 | 26 |
| No. of patients | 867 |  |  |  | 936 |  |  |  |
| Spearman rank correlation in cell-HR & patient-HR | 0.45 | 0.52* | 0.6* | 0.7* | 0.42* | 0.36 | 0.63 | 0.15 |

We also checked the Spearman rank correlation in high and low similarity thresholds between cell-HR and patient-HR with 43 frequently mutated cancer driver genes in breast cancer that are listed in COSMIC database. In higher similarity threshold, the correlation was higher (0.54), whereas in low similarity threshold the correlation was very low (0.015) (Fig 5). This shows that CTDPathSim correctly computed cell lines with high scores that were clinical biomarkers for patients’ hazard whereas low scores cell lines were not.

**Figure 5:**
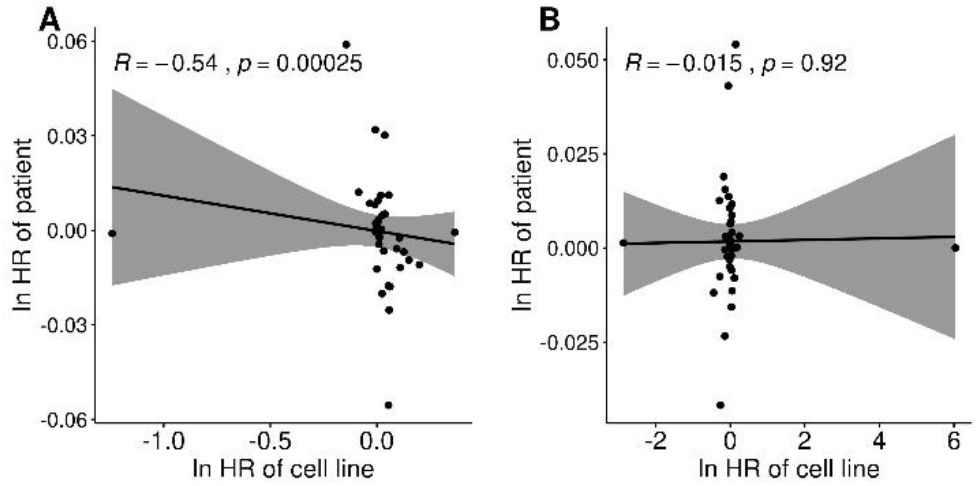
Scatter plots of cell lines’ hazard ratios (cell-HR) vs BRCA patients’ hazard ratios (patient-HR) with natural logarithm using 43 breast cancer driver genes. A) shows Spearman rank correlation with significant p-value in highly similar sample-cell lines group; B) shows Spearman rank correlation with insignificant p-value in lowly similar sample-cell lines group.

### 4.4. Aligned sample-cell line pairs had higher number of significant overlapping biological pathways than unaligned sample-cell line pair

We assessed CTDPathSim by comparing the overlapping biological pathways in highly similar sample-cell line pairs vs lowly similar sample-cell line pairs with TCGA BRCA samples and CCLE cell lines. We used biological pathways listed in KEGG [54] database for this analysis. For each cell line, we used frequently mutated breast cancer driver genes (listed in COSMIC) that were present in cell line’s DE genes and inferred KEGG pathways with significant enrichment (FDR corrected p-value < 0.01) in this gene list using R package *Pathfinder* [48]. Similarly, for each sample, we used breast cancer driver genes that were present in sample’s DE genes and inferred KEGG pathways with significant enrichment. We computed overlapping pathways for each cell line and sample pair with a hypergeometric p-value. We compared the median of these p-values in highly correlated sample-cell line pairs *i.e*. in aligned group vs lowly correlated sample-cell line pairs *i.e.* in unaligned group using the Wilcoxon rank-sum test. Interestingly, aligned group had lower p-values *i.e.* significant p-values (< 0.05), whereas, unaligned group had higher number of insignificant p-values (> 0.05) (Fig 6A). Also, we compared the median of overlapping pathways filtered by significant hypergeometric p values (< 0.01) between aligned group and unaligned group using the Wilcoxon rank-sum test. Aligned group was found to have more overlapping pathways than unaligned group with significant Wilcoxon p-value (Fig 6B). In Table 3, we listed the frequently appeared significantly overlapping KEGG pathways in aligned sample-cell line pairs within top 10 percentile. Interestingly, all the pathways in this list are cancer related pathways.

**Figure 6:**
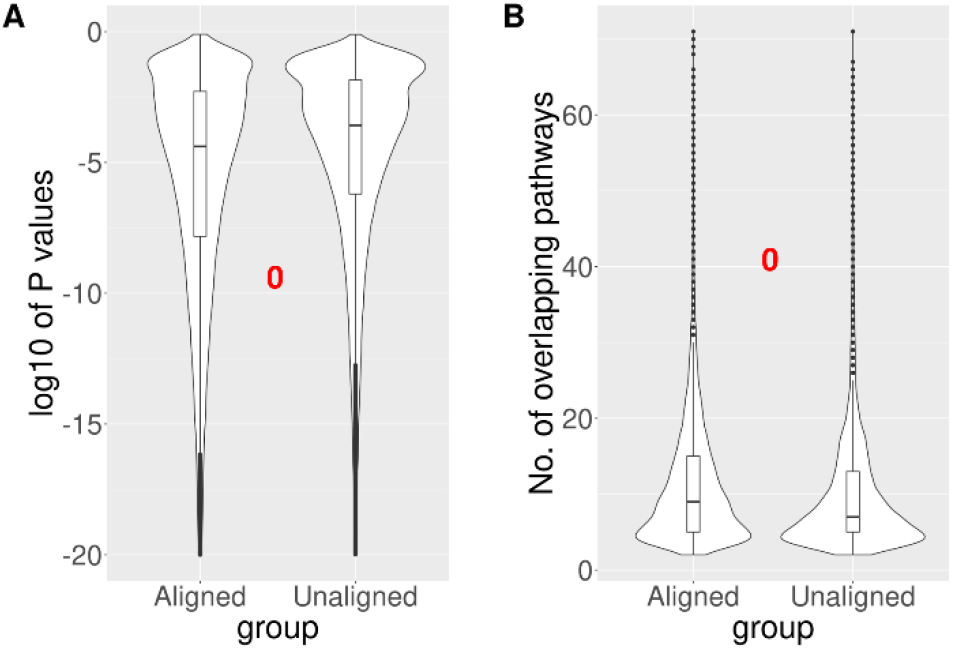
Overlapping KEGG pathways in highly similar sample-cell line pairs (aligned group) vs lowly similar sample-cell line pairs (unaligned group) with CTDPathSim’s computed scores between BRCA cohort and CCLE cell lines. **A)** logarithm of p-values of overlapping pathways in aligned group vs unaligned group. Aligned group shows lower median i.e. significant p-values compared to unaligned group; **B)** number of significant overlapping pathways in aligned vs unaligned group. Aligned group shows higher median value than unaligned group. Both the violin plots have significant Wilcoxson p-value (< 0.05) (in red).

**Table 3:**
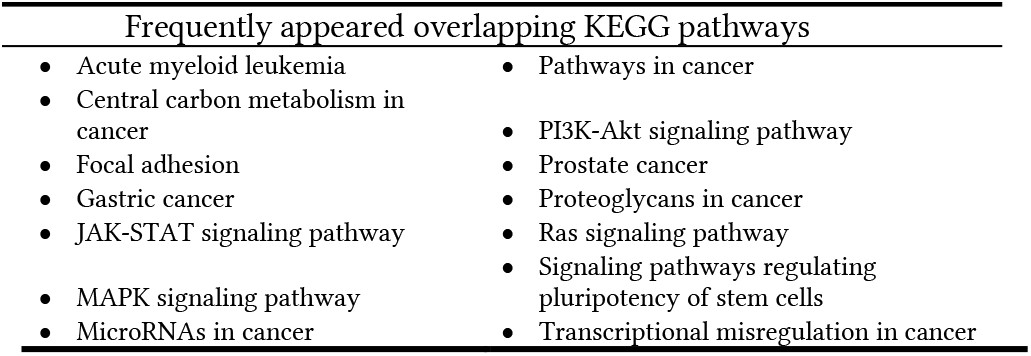
Frequently appeared overlapping KEGG pathways in highly similar sample-cell line pairs of BRCA sample and CCLE cell lines. These are the pathways that appear within top 10 percentile in frequently overlapping pathways

## 5. Discussions

We developed a computational pipeline, CTDPathSim, that utilized a pathway activity-based approach to compute similarity between primary tumors and cell lines at genetic, genomic, and epigenetic levels along with addressing bulk tumor heterogeneity.

We tested CTDPathSim on BRCA and OV samples from TCGA and cancer cell lines from CCLE. We computed the rank of 51 CCLE breast cancer cell lines using a median similarity score computed by CTDPathSim, between each cell line and all BRCA samples. Our top five breast cancer cell lines were, HMEL, HCC38, MDAMB453, CAL148 and HCC1599 (Supplemental Fig. 9). Similarly, we computed the rank of 47 CCLE ovarian cancer cell lines using a median similarity score each cell line and all OV samples (Supplemental Fig. 10). Our top five ovarian cancer cell lines were, KURAMOCHI, JHOS4, OELE, 59M and CAOV3. We also checked the rank of all cell lines of different cancer types, with a median similarity score to all samples in BRCA and OV cohorts. For BRCA, the top rank cell lines were the breast cancer cell lines (Fig. 7A) and for OV, the top rank cell lines were the ovarian cancer cell lines (Fig. 7B) among all the other cell lines from different cancer lineages.

**Figure 7:**
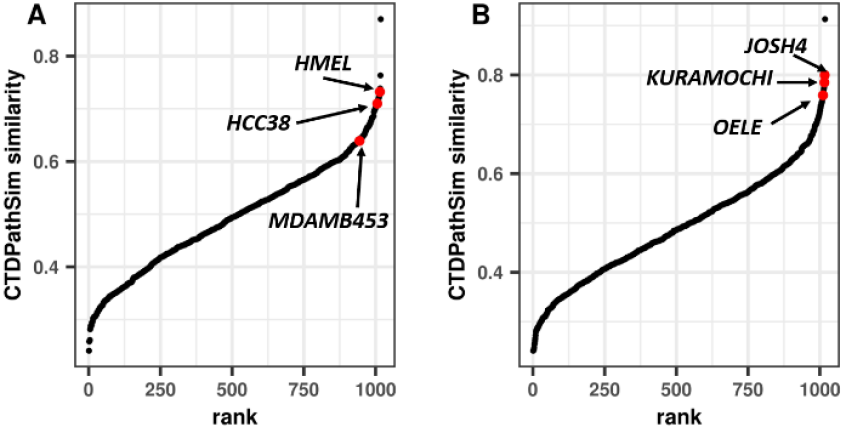
Rank of all 1,018 CCLE cell lines based on median similarity scores with all samples predicted by CTDPathSim. **A)** rank of all cancer cell lines in BRCA cohort with top rank breast cancer cell lines with red dots; **B)** rank of all cancer cell lines in OV cohort with top rank ovarian cancer cell lines with red dots.

Studies reported, MDAMB453, CAL148 and HCC1599 cell lines among top cell lines for resembling breast cancer samples [20], [49]. In an investigation of functional utility of ovarian cancer cell lines in pre-clinical trial, Coscia *et al.* [50] reported KURAMOCHI as one of the closest resemblance to the tumor samples. A recent study, using deep single cell RNA-Seq data, established this cell line as potential model of ovarian cancer subtype *in vitro* [51]. Domcke *et al.* [52] reported KURAMOCHI as one of the top cell lines for ovarian tumors. The same study reported CAOV3 as one of the poor choice to model ovarian tumors, however, in a xenograft-based study, Hernandez *et al.* [53] reported CAOV3 as one of the most resembling cell lines for ovarian cancer. Hence, our method was able to pick the top rank cell lines that were reported in existing *in vitro* studies.

CTDPathSim computed the sample-cell line similarity scores that were able to recapitulate the responses of known cancer drugs between cell lines and TCGA BRCA samples efficiently, compared to the state-of-the-art methods, namely, TSI method, SRCCM, TC analysis and Celligner (See Section 4.2). For CTDPathSim, majority of the drugs showed mismatch pattern in drug responses between cell lines and samples in lower similarities, whereas, there was significant trend in matching drug responses as the similarity scores were getting higher.

There were few drugs, such as Lapatinib and Doxorubicin, that did not show any clear pattern of drug response (Supplemental Fig. 3). We found that, the known drug response information for these drugs on cell lines were highly imbalanced having most of the cell lines as the resistant cell lines. Hence, to further increase the known drug response information between cell lines and TCGA breast cancer, we computed structural similarity scores between different drugs (see Materials & methods *2.3.2*). We used 17 highly similar (correlation ≥ 0.95) TCGA drugs to Lapatinib and treated the response of these drugs as proxy of Lapatinib’s response. For these 17 drugs, we found the structurally similar drugs in CCLE and GDSC databases. We tested all possible combination of these drug pairs between cell lines and samples. We used three TCGA drugs, namely Mitoxantrone, Pemetrexed and Fulvestrant, that passed our evaluation criteria (See Section 4.2) to be tested on behalf of Lapatinib’s response.

We observed that Mitoxantrone and Pemetrexed, can recapitulate the similar drug response pattern in high similarity scores of CTDPathSim, and showed the opposite trend in low similarities. On the other hand, for Fulvestrant, the drug response trend was not definitive (Supplemental Fig. 11).

In a lung cancer study, Ramlau *et al.* [54] showed that Lapatinib and Pemetrexed had a similar response rate in HER2-targeted therapies. A recent study suggested that Lapatinib might be a suitable treatment option for HER2-positive metastatic breast cancer patients that have become resistant to Trastuzumab *i.e.* Pemetrexed [55]. Also, Lapatinib, Mitoxantrone and Trastuzumab were found to be related with similar toxic effects in breast cancer patients [56],[57]. These studies point towards a correlation in drug responses between Lapatinib, Pemetrexed and Mitoxantrone that we also found in our study.

We also found, Doxorubicin as structurally similar drug with Lapatinib and hypothesize that their drug response pattern should be closely related. Interestingly, several studies established the efficacy of therapeutic outcomes when Doxorubicin used to be applied alone and with Lapatinib in Her2 positive patients [58] [59].

In conclusion, in this study, our goal was to compute a similarity score that connects a single cell line to a single patient sample. We evaluated the drug response concordance between our predicted score and the response of known drug therapies for sample-cell line pairs. Also, we identified lineage specific enrichment between samples cohort and cell lines. In the future versions of CTDPathSim, copy number and mutation data can be incorporated in computing similarity scores. We prepared a reference blood-derived methylation profiles for deconvolution that is applicable to pan-cancer [27, 30–33]. Hence, this study can be extended with wide range of pan-cancer cohort and can be evaluated based on many different cancer lineages. CTDPathSim is available with source codes and datasets at https://github.com/bozdaglab/CTDPathSim *under Creative Commons Attribution-NonCommercial 4.0 International Public License.*

## ACKNOWLEDGEMENTS

This work was supported by the National Institute OF General Medical Sciences of the National Institutes of Health under Award Number R35GM133657

